# Manufacturing-Aware Generative Model Architectures Enable Biological Sequence Design and Synthesis at Petascale

**DOI:** 10.1101/2024.09.13.612900

**Authors:** Eli N. Weinstein, Mattia G. Gollub, Andrei Slabodkin, Cameron L. Gardner, Kerry Dobbs, Xiao-Bing Cui, Alan N. Amin, George M. Church, Elizabeth B. Wood

## Abstract

We introduce a method to reduce the cost of synthesizing proteins and other biological sequences designed by a generative model by as much as a trillion-fold. In particular, we make our generative models manufacturing-aware, such that model-designed sequences can be efficiently synthesized in the real world with extreme parallelism. We demonstrate by training and synthesizing samples from generative models of antibodies, T cell antigens and DNA polymerases. For example, we train a manufacturing-aware generative model on 300 million observed human antibodies and synthesize *∼*10^17^ generated designs from the model, achieving a sample quality comparable to a state-of-the-art protein language model, at a cost of 10^3^ dollars. Using previous methods, synthesis of a library of the same accuracy and size would cost roughly a quadrillion (10^15^) dollars.

---

Generative models have achieved dramatic successes in designing novel and functional biological sequences. ^23;26;15;8;30^ Trained on natural data, these models learn the underlying constraints imposed by evolution and capture the range of outstanding biological possibilities. ^6;32^ Sampling from the model produces a diverse set of realistic designs. However, downstream testing of these designs is severely constrained by the cost of building them. This bottleneck limits our ability to discover sequences with desired properties, and limits our ability to improve and refine our machine learning models.

The conventional approach to synthesizing generated designs is to draw samples computationally and then synthesize each of these designs individually. ^23;26;15;8;30^ While generative models can produce astronomical numbers of novel sequences, and high-throughput experimental assays can evaluate billions of candidates or more, exact synthesis of individual sequences is limited by cost, and as a result most libraries do not exceed 10^5^ candidates in practice. ^10^ Degenerate codon methods can produce larger libraries, but they are uniformly random, with no connection to a given generative model.

Previously, in theoretical work, we introduced variational synthesis, a novel procedure for building generative biological sequence models and synthesizing samples from those models. ^31^ In this work, we implement variational synthesis in practice, at industrial scale, demonstrating successful manufacturing of about 10-100 quadrillion (10^16^-10^17^) samples from generative models in the real world across several important applications in protein design.

## Building Manufacturing-Aware Generative Biological Sequence Models

In this study, we develop a new approach to building generative biological sequence models, which allows their output to be manufactured physically at large scale. The key idea is to integrate knowledge of DNA synthesis into the model architecture. Such manufacturing-aware generative models can produce designs that are just as diverse and realistic as those of other modern generative models. However, their designs can also be synthesized *in vitro* in extreme parallel. We refer to such models as *variational synthesis models*. ^31^

Formally, a variational synthesis model describes a distribution p_*θ*_(*x*) over amino acid sequences *x* controlled by parameters *θ*, such that each element of *θ* corresponds to an experimentally controlled parameter in a DNA synthesis protocol. In particular, the distribution p_*θ*_(*x*) is the distribution of amino acid sequences produced in the laboratory by synthesis reactions with the experimental parameters *θ*. We can train variational synthesis models on data by adjusting the parameters *θ* to find an optimal *θ*^*⋆*^, such that p_*θ*_^*⋆*^ (*x*) closely approximates the data distribution. Then, samples from the trained model p_*θ*_^*⋆*^ (*x*) can be manufactured in the physical world by running synthesis reactions with the learned parameters *θ*^*⋆*^.

To put it otherwise, the overall workflow of variational synthesis is as follows (Figure 1a). We begin with a variational synthesis model, with a manufacturing-aware architecture suited to the experimental synthesis platform available. We train this model on a large dataset of biological sequence examples. Then, we can sample designs from this generative model not only *in silico* but also *in vitro*, by running DNA synthesis reactions according to the learned parameters of the model. The result is a very large number of manufactured samples from the generative model.

**Figure 1:**
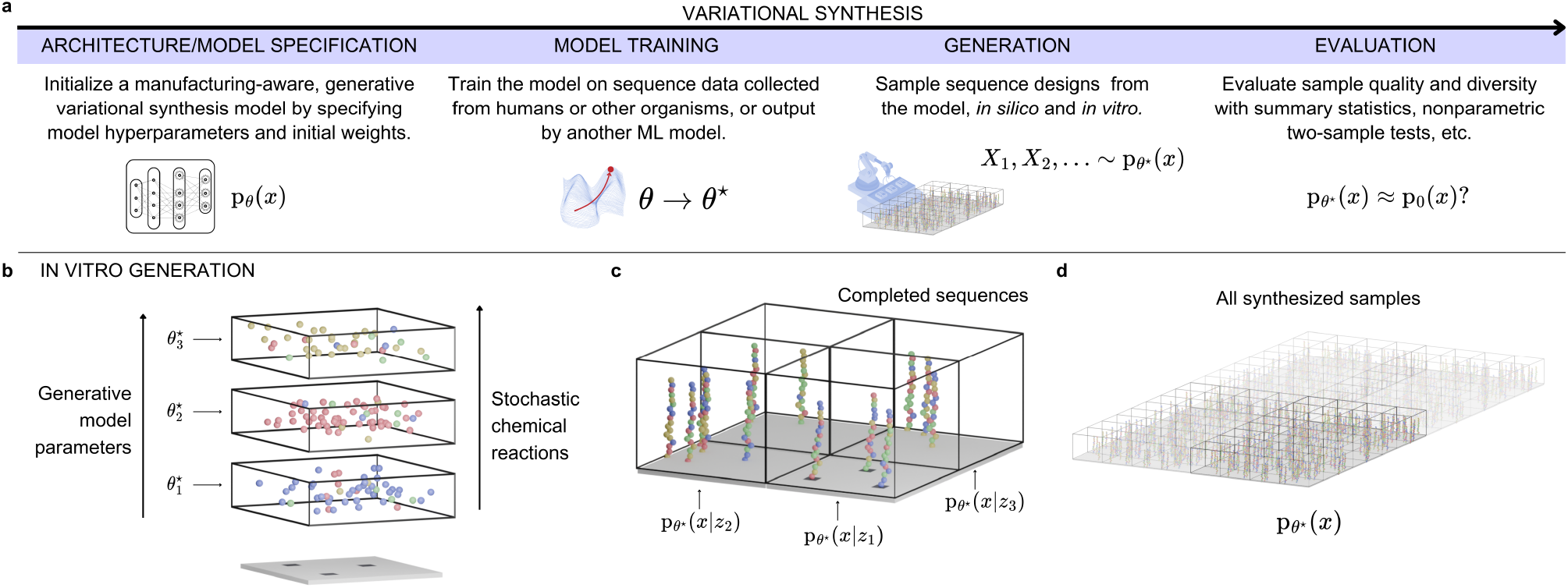
Variational synthesis models control DNA synthesis reactions to manufacture designs. **a**, Overview of the variational synthesis workflow. Here, p_*θ*_^*⋆*^ (*x*) is the variational synthesis model and p_0_(*x*) is the data distribution that the model is learning. **b-d**, Details of the *in vitro* generation process. **b**, Each parameter of the trained variational synthesis model 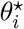 determines an experimental parameter in a stochastic chemical reaction. To synthesize complete sequence designs, we run a series of such controlled reactions. **c**, We repeat the process across different reaction compartments, with each compartment corresponding to a different latent variable *z*_*i*_ in the generative model. **d**, The end result, using many reaction compartments, is synthesized samples from the generative model. Each molecule of DNA is an independent sample *X ∼* p_*θ*_^*⋆*^ (*x*). The total number of molecules/samples reaches *∼*10^16^-10^17^.

With a conventional generative model, the only available way to synthesize designs is to sample designs on the computer, and then synthesize each design individually. With a variational synthesis model, however, computational sampling and chemical synthesis are more deeply integrated. This is because the parameters of a variational synthesis model are not simply uninterpretable weights and biases. Rather, each *in silico* parameter of the model maps to a specific experimental parameter of a DNA synthesis procedure, such as the concentration of a certain reagent, or the timing of a particular reaction (Figure 1b-d). A variational synthesis model thus provides a highly specific experimental protocol, with step-by-step instructions for large-scale laboratory synthesis of its designs.

Variational synthesis models enable synthesis of extremely large numbers of model samples via the controlled use of stochastic chemical reactions. All generative models depend on controlled randomness to produce diverse designs. Conventionally, this randomness is introduced by a random number generator on the computer. Variational synthesis models can instead exploit the physical randomness inherent in stochastic chemical reactions. As the synthesis process proceeds, each molecule of DNA randomly encounters a different series of reactants. As a result, every DNA molecule corresponds to a new design, an independent sample from the model. So if the synthesis reactions yield a picomole of DNA, we will have synthesized six hundred billion samples from the generative model. Despite the randomness, however, the population of DNA molecules as a whole remains highly controlled, as the experimental parameters of each stochastic reaction are set by the learned parameters of the generative model.

We previously designed and computationally evaluated several variational synthesis models with different model architectures based on different stochastic synthesis technologies. ^31^ Following these evaluations we decided to focus on an oligosynthesis technology that delivers mixtures of bases in each round of nucleotide addition (Appendix B). Working with a large commercial DNA synthesis provider, we developed both (a) customized synthesis protocols and (b) variational synthesis models describing the distribution of DNA and amino acid sequences produced by these specific protocols. Our variational synthesis models account for complex manufacturing constraints, and the model-designed reactions have been carried out repeatedly at scale with automation. With appropriate calibration and quality control, we found that we could reliably produce samples from a specified variational synthesis model with 97% accuracy per base (Table S1, Figure S1). We achieve yields on the order of *>*10 nanomoles of full-length DNA, corresponding to *>*10^16^ model samples (petascale synthesis).

We developed optimization procedures that allow our synthesis models to be trained on datasets with hundreds of millions of sequences. As with other generative models, we use GPU-accelerated stochastic optimization with minibatches of data. However, many experimental parameters in synthesis reactions are discrete and hence do not admit backpropagation. We therefore combine conventional stochastic gradient descent methods with stochastic discrete optimization methods, based on adaptive expectation maximization. ^31^

## Synthesizing a Pan-human Antibody Repertoire

Antibodies are central to the human adaptive immune system and a key source of new medicines, including monoclonal antibody therapies, antibody-drug conjugates, and cell therapies.^14;9^ Large scale antibody synthesis allows for detailed experimental study of antibodies’ diverse properties, enabling characterization of human immunity at the population level, and discovery of novel therapeutic candidates. ^33^ We set out to design and build antibody sequences spanning the full range of human post-selection antibody diversity (Figure 2a). To accomplish this, we first trained a variational synthesis model on a large and diverse dataset of post-selection human antibodies. We then used the model to generate CDRH3 samples of diverse lengths, *in silico* and *in vitro*. We validated the model’s *in silico* performance by checking how well the distribution of samples matched held-out data. We validated its *in vitro* performance by sequencing a subsample of manufactured designs and similarly evaluating them against held-out data. We found that our generative model performs well *in silico*, accurately predicting held-out sequences and generating realistic novel sequences. We further found that our generative model performs well *in vitro*, manufacturing a petascale library that closely resembles real, unseen human antibody CDRH3s – as much as do *in silico* designs from an *in silico*-only protein language model.

**Figure 2:**
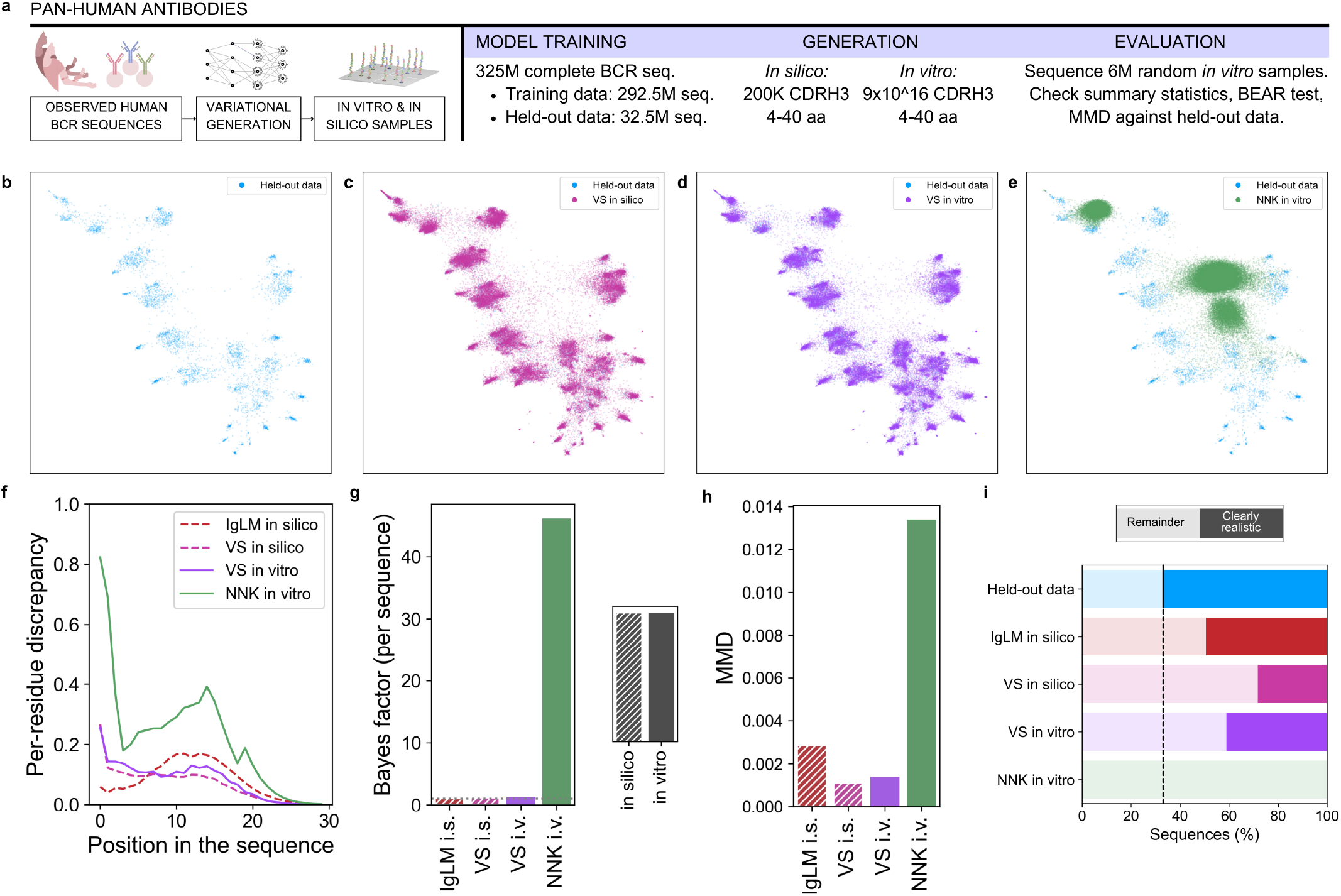
Synthesizing samples from a generative model of the pan-human antibody repertoire. **a**, Overview of the workflow. The data consists of antibody CDRH3 sequences collected from eleven thousand human repertoires. We train a variational synthesis model on this data, generate samples from the model *in silico* and *in vitro*, and evaluate sample quality. **b**, Low-dimensional representation of sequences from the held-out data. **c**, Low-dimensional representation of sequences from the held-out data, together with sequences generated *in silico* by the variational synthesis model. **d**, Low-dimensional representation of sequences from the held-out data, together with sequences manufactured *in vitro* by variational synthesis. **e**, Low-dimensional representation of sequences from the held-out data, together with sequences manufactured *in vitro* using degenerate codon synthesis (NNK codons). **f**, Per-residue discrepancy between generated sequences (*in silico* or *in vitro*) and the held-out data. The discrepancy is based on total variation distance (Appendix C.1). **g**, Per-sequence Bayes factor of the BEAR two-sample test, comparing generated sequences (*in silico*: i.s. or *in vitro*: i.v.) and the held-out data. The dashed line indicates a Bayes factor of one; values lower than one favor the hypothesis that the model and data distributions are identical. **h**, Estimated maximum mean discrepancy (MMD) between generated sequences and the held-out data. An MMD of zero indicates a perfect distribution match. **i**, Fraction of generated sequences that are considered *clearly realistic* based on a high MMD witness function score. Dashed line indicates the percentile threshold set to define *clearly realistic* based on the held-out data.

To begin, we compiled a training dataset of 325 million unique human heavy-chain antibody sequences, drawn from a diverse set of 11,271 repertoires in the OAS database (Appendix D). ^19^ We excluded patients with autoimmune disorders and related conditions, to ensure the training data consists of functional and non-autoreactive sequences. Trained on heavy-chain CDR3 sequences, the variational synthesis model estimates the distribution underlying all healthy human antibody CDRH3 sequences, covering both seen and unseen sequences. The model achieves an average per-residue perplexity on a held-out test set of 10.6, comparable to current protein language models and other deep generative antibody models. ^20^ Training set perplexity is similar to the measured test set perplexity of 10.6, indicating strong generalization from seen to unseen antibodies.

We first sought to evaluate the *in silico* sample quality of the variational synthesis model, i.e. its performance in the absence of any manufacturing errors. We began by comparing summary statistics of the generated samples to held-out data. We found the variational synthesis model accurately captures the average amino acid usage across positions (Figure 2f, Appendix C.1). Lowdimensional representations suggest qualitatively that samples from the variational synthesis model closely follow the distribution of the held-out data (Figure 2bc, Appendix C.2).

To more stringently evaluate sample quality, we used nonparametric two-sample test evaluations based on the Bayesian embedded autoregressive (BEAR) model and the maximum mean discrepancy (MMD) with a biological sequence kernel. ^1;4;3^ These recently-developed methods rigorously check the entire distribution of model samples for mismatch with the held-out data, rather than just checking a finite number of metrics. They thus help avoid the possibility that the generative model performs well according to hand-picked scores, but poorly along other dimensions. The two evaluations are complementary, with BEAR more focused on short-range correlations (it relies on variable-length kmer statistics) and MMD focused on more global structure (it relies on padded Hamming distances between sequences) (Appendix C.3). We compared the variational synthesis model to a state-of-the-art antibody language model, IgLM. ^27^ IgLM was trained on a very similar dataset of human antibodies, also based on OAS (Appendix D.1). It produces realistic human antibody sequences *in silico*, but lacks a manufacturing-aware architecture, so *in vitro* samples must be synthesized individually.

We find that *in silico* samples from our variational synthesis model achieve a close match to held-out data, with sample quality similar to IgLM (Figure 2gh, Appendix D.1). In detail, the BEAR test provides a Bayes factor describing the odds in favor of the model and data distributions being different. We report the per-sequence geometric average Bayes factor, which tells us the evidence contributed by each sequence (in units of odds). Applied to our data, the BEAR test says that both generative models achieve a close though not perfect match to the data, with an average Bayes factor close to one: 1.31 for the variational synthesis model and 1.02 for IgLM (Figure 2g). The MMD quantifies the mismatch between model samples and held-out data in terms of the worst-case difference in their average phenotype (Appendix C.3). ^32;2^ The MMD confirms that variational synthesis model samples are highly realistic, and indeed says the variational synthesis model outperforms IgLM (Figure 2h). Overall, these evaluations suggest that variational synthesis models are powerful generative models that can achieve comparable *in silico* performance to state-of-the-art deep generative models.

We further sought to quantify the fraction of *in silico* model samples that are highly plausible human antibodies. To do so we used the witness function of the MMD, which scores how much a sequence looks like real data as opposed to a sample from the model. ^13^ Low scores suggest a sequence is an outlier and might not have the functional properties of a real human antibody. To be conservative, we consider a model-generated sequence to be *clearly realistic* only if its score is in the top 67% of witness function scores for real, held-out human antibody sequences (Appendix C.4). By this metric, 28% of samples from the variational synthesis model are clearly realistic, versus 49% of samples from IgLM (Figure 2i, Figure S5).

Next, we manufactured CDRH3 samples of length 4-40 amino acids from our trained variational synthesis model. We achieved a yield of roughly 150 nanomoles, 9 × 10^16^ molecules. We sequenced a random subset of these samples: six million after paired-end read assembly and filtering (Appendix A). We find the manufactured variational synthesis library has only a slight degradation in performance compared to our *in silico* predictions, with the manufactured samples maintaining similar average amino acid usage (Figure 2f) and similar low-dimensional representations to real held-out data (Figure 2d, Figure S2a, Figure S3a). As an additional comparison, we also constructed a degenerate codon library, based on NNK codons (where N is randomly A, C, G or T and K is randomly G or T; Appendix C.5). Such NNK libraries can achieve similar sequence diversity to variational synthesis libraries, as they also rely on stochastic synthesis reactions, but they are uniformly random with no relationship to a trained generative model. We found that sequences in the manufactured NNK library show dramatically different amino acid usage and low-dimensional representations compared to samples from the variational synthesis model (Figure 2ef, Figure S2d, Figure S3d).

The BEAR and MMD tests show the manufactured variational synthesis samples achieve a close match to held-out data, only slightly worse than the *in silico* samples from the variational synthesis model (Figure 2gh). For example, the BEAR test Bayes factor increases from 1.31 to 1.39. Overall, the quality of the *in vitro*, manufactured samples is not only substantially better than that of NNK libraries, but is even comparable to the quality of *in silico* samples from IgLM: *in vitro* samples from the variational synthesis model have lower MMD than *in silico* samples drawn from IgLM (Figure 2h).

According to the witness function, 39% of the manufactured sequences are clearly realistic antibody sequences, compared to IgLM’s *in silico* fraction of 49% (Figure 2i). By contrast, in the NNK library, only 0.08% of sequences are clearly realistic.

In summary, variational synthesis models are able to create tens of quadrillions of samples from the pan-human antibody repertoire distribution, with *in vitro* sample quality similar to that of an *in silico*-only, state-of-the-art deep generative antibody model. We conservatively estimate that the variational synthesis library contains 10^16^ realistic antibody sequences, which we made at a cost of roughly $10^3^. By comparison, conventional oligosynthesis approaches cost roughly $0.01 per base, so a library of the same size and lengths, consisting of samples from a conventional generative model, would cost roughly $10^15^. We therefore achieve an estimated cost reduction of around a trillion-fold.

## Synthesizing the HLA-A*02:01 Epitope Repertoire

Peptide epitopes presented on HLA molecules are recognized by human T cells, leading to an adaptive immune response to pathogens and tumors. These linear epitopes form the basis for T cell vaccines against infectious disease and cancer. Large scale synthesis of epitopes allows for experimental determination of the antigens that human T cells respond to, and discovery of novel vaccine candidates. ^11;7^ We set out to comprehensively explore the space of T cell antigens, and synthesize the full range of peptides that bind a common HLA allele, HLA-A*02:01. To do so, we trained a variational synthesis model on a dataset of peptides predicted to bind HLA-A*02:01, then manufactured *∼*10^16^ samples from the variational synthesis model and validated the library by sequencing (Figure 3a). We find a close correspondence between the distribution of synthesized sequences and the distribution of predicted HLA-A*02:01 binders.

**Figure 3:**
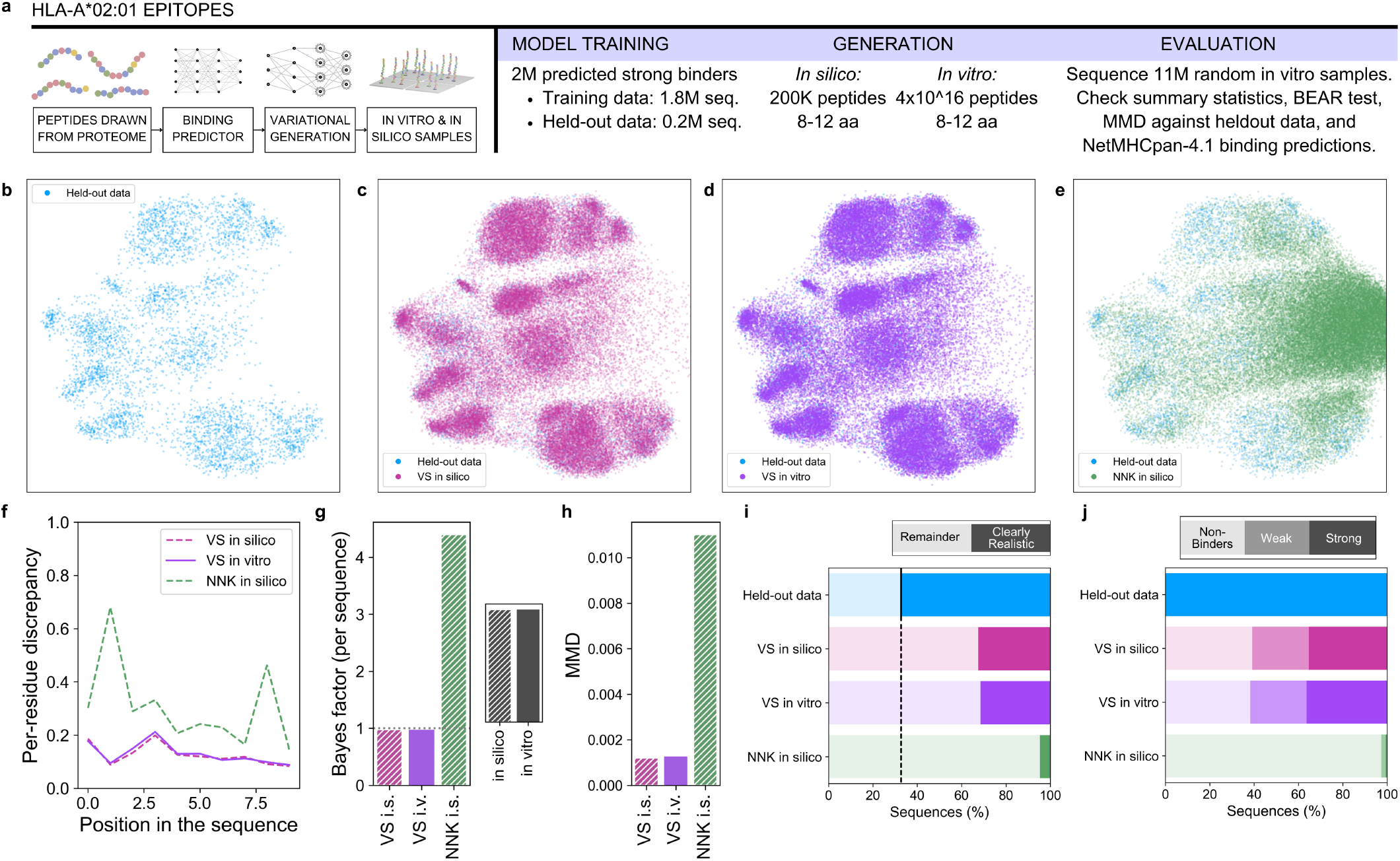
Synthesizing samples from a generative model of antigens presented on HLA-A*02:01. **a**, Overview of the workflow. The training dataset consists of diverse antigens (human and non-human) predicted to bind HLA-A*02:01. **b**, Low-dimensional representation of sequences from the held-out data. **c**, Low-dimensional representation of sequences from the held-out data, together with sequences generated *in silico* by the variational synthesis model. **d**, Low-dimensional representation of sequences from the held-out data, together with sequences manufactured *in vitro* by variational synthesis. **e**, Low-dimensional representation of sequences from the held-out data set, together with sequences generated *in silico* with NNK codons. **f**, Per-residue discrepancy between generated sequences (*in silico* or *in vitro*) and the held-out data. The discrepancy is based on total variation distance (Appendix C.1). **g**, Per-sequence Bayes factor of the BEAR two-sample test, comparing generated sequences (*in silico* or *in vitro*) and the held-out data. **h**, Maximum mean discrepancy (MMD) between generated sequences and the held-out data. **i**, Fraction of generated sequences that are considered *clearly realistic* based on a high MMD witness function score. **j**, Fraction of sequences classified as non-, weak, or strong binders by NetMHCpan-4.1.

First, we constructed a large dataset of natural peptides predicted to bind HLA-A*02:01, covering both human (neo)antigens and pathogen antigens (Appendix E). We began by sampling *in silico* peptides according to the natural amino acid frequencies and length range, 8-12 amino acids. Then, we virtually screened these peptides for binding against HLA-A*02:01, using NetMHCpan, a state-of-the-art machine learning model that returns a prediction of binding strength for a given HLA-epitope pair. ^22^ The resulting dataset consists of 2 million samples from the conditional distribution of natural peptides predicted to bind HLA-A*02:01. We trained a variational synthesis model on 90% of this dataset, achieving a perplexity of 14.2 per amino acid on the held-out 10% of the data, versus 20.6 for NNK libraries (Appendix C.5). Since the data is generated synthetically, we can compute the true data perplexity as 11.4, modestly lower than that achieved by the synthesis model.

We synthesized samples of epitopes with lengths 8-12 amino acids, according to the parameters of the variational synthesis model. We achieved a yield of roughly 75 nanomoles, corresponding to 4 × 10^16^ molecules. We then verified the samples by sequencing a subset (11 million reads after assembly and filtering) and applying summary statistic and nonparametric two-sample evaluations. We find close correspondence between the synthesized DNA and held-out data, in average amino acid usage and in low-dimensional representations (Figure 3df, Figure S2b, Figure S3b). In particular, the manufactured variational synthesis libraries show dramatic improvements over NNK libraries, based on *in silico* sampled NNK sequences (Figure 3ef, Figure S2e, Figure S3e, Appendix C.5). Nonparametric two-sample test evaluations also show close correspondence between synthesized DNA and held-out data, again with large improvements over NNK libraries (Figure 3gh). Indeed, the BEAR test suggests the synthesized sequences are statistically indistinguishable from held-out data: the Bayes factor is below one, favoring the hypothesis that the model and data distributions are the same. According to the witness function score, 31% of the sequences manufactured by variational synthesis are clearly realistic, versus 4.6% of NNK library sequences (Figure 3i, Figure S6a). For this library, we can also check the quality of our manufactured samples using the binding predictor. We find that 36% of the sequences manufactured *in vitro* by variational synthesis are predicted to strongly bind HLA-A*02:01 by NetMHCpan, with an additional 25% predicted to bind weakly (Figure 3j, Table S2). By contrast, 0.7% of NNK library sequences are predicted to bind strongly, and 2% weakly.

Overall, then, variational synthesis enables accurate manufacturing of about 1 × 10^16^ samples from the global distribution of T cell epitopes predicted to present strongly on HLA-A*02:01, in a library where half of all sequences are predicted to be either a strong or weak binder.

## Synthesizing DNA Polymerases Across Evolution

DNA polymerases play an essential role in modern biotechnology, with major applications including polymerase chain reaction (PCR) and next generation sequencing (sequencing by synthesis). Large scale synthesis of polymerase variants enables screening for enzymes with greater fidelity, higher thermostability, non-natural substrates, and other desirable properties. ^18^ DNA polymerases are ancient and found across life, so we set out to synthesize novel DNA polymerases spanning the range of evolutionary possibility. We trained a variational synthesis model on a dataset of DNA polymerase sequences predicted to be generated by evolution, then manufactured *∼*10^16^ samples from the variational synthesis model, validated the library by sequencing, and confirmed that the distribution of manufactured sequences corresponds closely to held-out data (Figure 4a).

**Figure 4:**
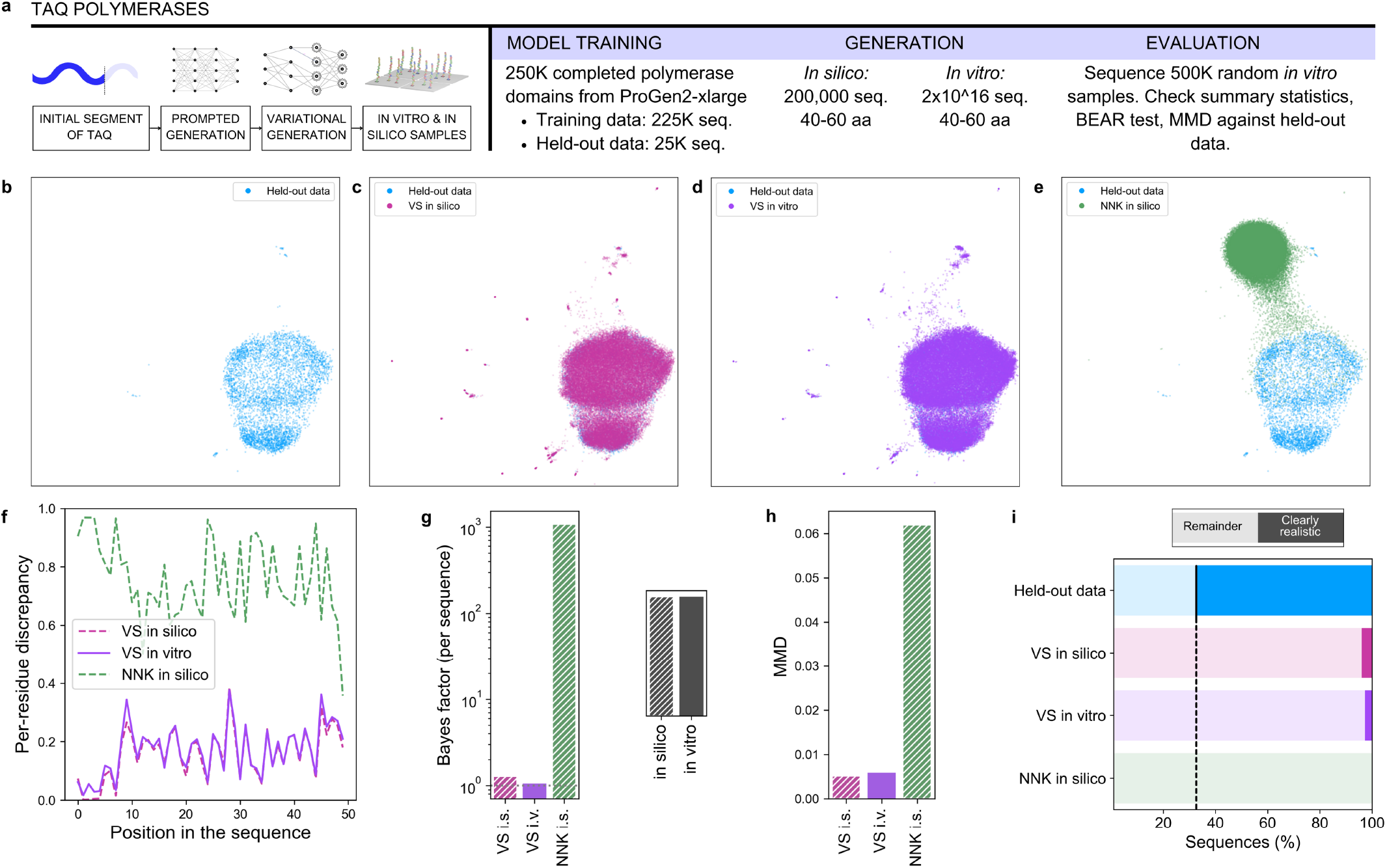
Synthesizing samples from a generative model of DNA polymerases. **a**, Overview of the workflow. The data consists of completed *Taq* polymerase sequences generated by an evolutionary protein language model. We train a variational synthesis model on this data, generate samples from the model *in silico* and *in vitro*, and evaluate sample quality. **b**, Low-dimensional representation of sequences from the held-out data. **c**, Low-dimensional representation of sequences from the held-out data, together with sequences generated *in silico* by the variational synthesis model. **d**, Low-dimensional representation of sequences from the held-out data, together with sequences manufactured *in vitro* by variational synthesis. **e**, Low-dimensional representation of sequences from the held-out data set, together with sequences generated *in silico* with NNK codons. **f**, Per-residue discrepancy between generated sequences (*in silico* or *in vitro*) and the held-out data. The discrepancy is based on total variation distance (Appendix C.1). **g**, Per-sequence Bayes factor of the BEAR two-sample test, comparing generated sequences (*in silico* or *in vitro*) and held-out data. **h**, Maximum mean discrepancy (MMD) between generated sequences and the held-out data. **i**, Fraction of generated sequences that are considered *clearly realistic* based on a high MMD witness function score.

We used a protein language model to provide an estimate of the distribution of DNA polymerases likely to be produced by evolution. In particular, we used ProGen2, which has been trained on protein data drawn from across evolution, and so provides a global estimate of sequences’ evolutionary fitness. ^17;32^ We conditioned (prompted) the model with an initial segment of the *Taq* polymerase, a thermostable DNA polymerase widely used in PCR. Specifically, we conditioned on all but the last 50 amino acids of *Taq*, then generated new sequences with lengths up to 60 amino acids for the remainder of the enzyme (Appendix F). Each generated design represents a predicted evolutionary possibility for a complete protein. We drew 250,000 samples from ProGen2, and used these samples as training data for a variational synthesis model. Trained on 90% of the data, the variational synthesis model achieves a perplexity of 3.54 per amino acid on the held-out 10% of the data, versus 20.44 for NNK libraries. Since the data is generated from a model with tractable likelihoods, we can estimate the true perplexity as 2.57, modestly lower than the variational synthesis model (Appendix F).

We synthesized samples of the final region of the *Taq* polymerase, with length 40-60, according to the parameters of the trained variational synthesis model. We achieved a yield of roughly 35 nanomoles, corresponding to 2 × 10^16^ molecules. We sequenced a random subset: 500,000 reads after assembly and filtering. The distribution of synthesized DNA and held-out data match closely, according to summary statistics and nonparametric two-sample evaluations (Figure 4dfgh, Figure S2c, Figure S3c). As elsewhere, we find large improvements over NNK libraries (Figure 4efgh, Figure S2f, Figure S3f). According to the witness function analysis, 2.8% of the sequences manufactured by variational synthesis are clearly realistic, while we could not find any clearly realistic sequences among 200,000 sampled from the NNK library (*<*0.0005% clearly realistic) (Figure 4i, Figure S6b).

In short, we have shown that variational synthesis models can be trained on data generated by a conventional, non-manufacturing-aware generative model, and the resulting quadrillion manufactured samples closely follow that model’s designs.

## Discussion

We have shown that variational synthesis models enable biological sequence design and real-world synthesis at petascale. Their *in vitro* sample quality, in terms of realism and diversity, is comparable to that of state-of-the-art generative models. They are broadly applicable across different protein families.

Variational synthesis models achieve petascale synthesis via a manufacturing-aware model architecture, which accounts for the possibilities and constraints of stochastic DNA synthesis technology. Variational synthesis models are therefore both enabled and limited by available experimental tools for DNA synthesis. For example, although we have focused on relatively short mixed-length protein regions, this is not a constraint of variational synthesis *per se* but rather of the underlying oligosynthesis technology chosen for our example models. It is possible to design variational synthesis models that incorporate knowledge of DNA assembly techniques, and other experimental strategies for constructing long DNA sequences. ^31^ Developing and using such models will be important for many application areas going forward. It may also be fruitful to develop novel DNA synthesis technologies that are specifically suited to variational synthesis applications.

Variational synthesis models can integrate information beyond just biological sequences. For example, this may include information about proteins’ functional properties, measured by experimentally testing *in vitro* samples from an earlier variational synthesis model. Or, this information may include domain-specific proxies of biological function, such as 3D protein structure. In this paper, we integrate auxiliary information by using auxiliary models (i.e. NetMHCpan and ProGen). First we condition and generate samples with the auxiliary model, then we train a variational synthesis model on those samples. However, training one generative model on data produced by another risks compounding errors. ^28^ Going forward, it will be important to develop improved methods for conditioning variational synthesis models on auxiliary information and design goals.

A persistent theme in machine learning is the fundamental importance of large, relevant data sets. This has been a key bottleneck for biology applications in particular. Variational synthesis libraries offer an ideal input for experimental assays that create new datasets. By training a variational synthesis model on real biological sequences, we can ensure the resulting data contains relevant, in-distribution examples. By manufacturing sequences at scale, we can ensure the datasets are very large and fully cover the space of sequences we wish to understand.

Of course, as limits on synthesis are removed, other challenges come to the fore in building massive datasets. Current ultra-large scale experimental assays are designed for screening libraries of unrealistic and uninformative sequences, such as produced by NNK libraries. As a consequence, they typically focus on coarse-grained measurements of protein function, and discard information about non-functional sequences entirely. In a variational synthesis library, sequences are realistic and relevant. Now, the outstanding challenge is to create experimental technologies – including systems for expression, display, screening, characterization, and sequencing – that can extract meaningful information about every sequence in the library. This problem grows even more acute when screening variational synthesis libraries against one another, e.g. to map interactions across binders and targets.

The future rewards of overcoming these challenges are great: with petascale libraries and experimental systems to explore them, biological datasets may quickly rival in size and quality the internet-scale datasets available for images and text. With such data, we can bring to bear the full armamentarium of modern machine learning techniques, to answer the important questions facing biology.

## Supplementary information

## A Sequencing protocol

### A.1 Sequencing Library Preparation

250ng of DNA were sampled from each library. The sequencing library was prepared using xGen™ssDNA & Low-Input DNA Library Preparation Kit (IDT) following manufacturer’s instructions with the exception that post-extension cleanup was performed using MinElute PCR Purification Kit (Qiagen). All other cleanups were performed using 1.8x SPRI bead ratio.

### A.2 Illumina Sequencing

Dual indexed library samples were quantified using Kapa Library Quantification Kit (Roche). Pooled library samples were sequenced in a 2×150bp configuration on a NextSeq 2000 using P1 Reagents, 300 cycle Kit (Illumina). The resulting paired reads were basecalled with `bcl2fastq` v2.20 and assembled with `PEAR` ^34^.

We obtained 5,706,832 sequences for the antibody variational synthesis library, 10,844,511 sequences for the epitope library, and 510,864 for the DNA polymerase library.

## B Variational synthesis implementation

A detailed description of variational synthesis can be found in Weinstein et al. (2022).^31^ Code implementing synthesis models can be found at https://github.com/debbiemarkslab/variational-synthesis. Details of the specific synthesis model architectures used in this paper were customized based on proprietary platform information owned by the commercial synthesis provider.

## C Evaluations

In this section we detail the methods used to evaluate designed and synthesized libraries.

### C.1 Average amino acid and nucleotide usage

#### C.1.1 Per-position error

We evaluated the match between real data and manufactured sequences in terms of average amino acid usage per position. For each position, we quantify the absolute error in amino acid frequency using an L1 discrepancy: the total variation distance between the marginal distribution over amino acids at each position. To account for length differences among sequences, we pad each sequence on the right with a stop character, $, and include this in our alphabet.

Formally, let *ℬ* denote the amino acid alphabet. Let p(*x*_*j*_) denote the marginal likelihood of a sequence distribution p(*x*) at position *j*, such that p(*x*_*j*_ = *b*) for *b ∈ ℬ* is the probability of amino acid *b* appearing at position *j*, and p(*x*_*j*_ = $) is the probability of a sequence being shorter than length *j*. Hence, Σ_*b∈ ℬ*∪{$}_ p(*x*_*j*_ = *b*) = 1. Now, the discrepancy in average amino acid usage at position *j*, between distributions p and q, is

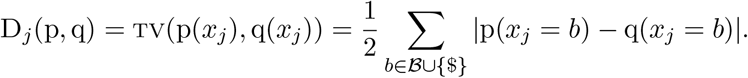

In practice, we estimate the per-position discrepancy from samples by taking the empirical frequency 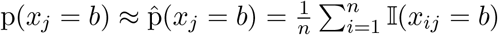 where *n* is the number of samples and 𝕀(*·*) is the indicator function that takes value one when the statement is true and zero otherwise. For Figure 2f, Figure 3f, and Figure 4f, we compare the data distribution to the manufactured variational synthesis distribution by evaluating 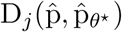, and similarly for the comparison methods. For *in silico* libraries (such as *in silico* variational synthesis models or IgLM) we use 200,000 samples.

**Table S1:** Manufacturing accuracy of variational synthesis models on different libraries. See Appendix C.1.2 for details of the accuracy calculation.

|  | Antibody | Epitope | DNA polymerase | Average |
| --- | --- | --- | --- | --- |
| Per-base (nucleotide) accuracy | 95.80% | 97.56% | 97.63% | 97.00% |
| Per-residue (amino acid) accuracy | 94.51% | 95.85% | 96.49% | 95.62% |

#### C.1.2 Overall accuracy

To compute an overall measure of accuracy, we average the discrepancy across positions. We then define the accuracy as one minus the error,

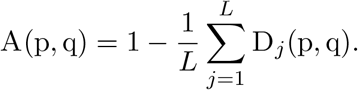

We estimate this empirically from samples as

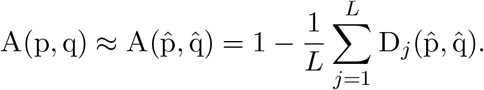

We set the maximum length *L* to the typical length of sequences in each dataset, rather than the maximum length, since our aim here is to evaluate our ability to accurately match amino acid or nucleotide usage, rather than length. In practice, the length distributions was unimodal in each dataset, and we set *L* to the mode: *L* = 15 for the antibody dataset, *L* = 9 for the epitope dataset, *L* = 49 for the DNA polymerase dataset. This is the notion of accuracy we report in Table S1.

This notion of accuracy appropriately generalizes the standard notion of accuracy used when the goal is to synthesize a specific sequence. For example, consider a situation where the goal is to synthesize a sequence *ACG*, but only 95% of the sequences we manufacture have an *A* at position one, 95% have a *C* at position two, and 95% have a *G* at position three. Then we would say the average accuracy is 95%. Mathematically, the goal of synthesizing *ACG* corresponds to a target distribution of p(*x*) = *δ*_*ACG*_(*x*), a delta distribution at *ACG*. This corresponds to a target marginal distribution of p(*x*_1_ = *A*) = 1 and p(*x*_1_ = *b*) = 0 for *b* ≠ *A*. Meanwhile, the realized marginal distribution at this position is 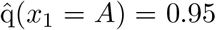 and 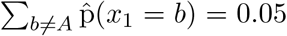. This gives 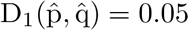, and, extending the same logic, 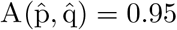. So, our accuracy metric reduces to the standard accuracy metric used when the goal is to synthesize a specific sequence.

### C.2 Low dimensional representations

To qualitatively evaluate the areas of sequence space covered by the target data and our libraries, we computed two-dimensional sequence representations. To do this we (i) computed a matrix of pairwise Hamming distances (with all sequences padded on the right to the same maximum length) between all sequences from the held-out data, the designed and manufactured variational synthesis libraries, and the NNK library, (ii) transformed this distance matrix into Gram matrix using the kernel described in Appendix C.3, and (iii) reduced the dimension of this matrix using UMAP. ^16^ We then showed only the held-out set (Figure 2b, Figure 3b), the held-out set overlapped with the variational synthesis library (Figure 2c, Figure 3c) and the held-out set overlapped with the NNK library (Figure 2d, Figure 3d), to illustrate how the much of the target data distribution is covered by each library. For the sake of visual interpretability, we use a subsample of 5,000 sequences from the held-out data and 50,000 from each library; we take more sequences from the libraries since the manufactured variational synthesis libraries are much larger than the held-out data set.

**Figure S1:**
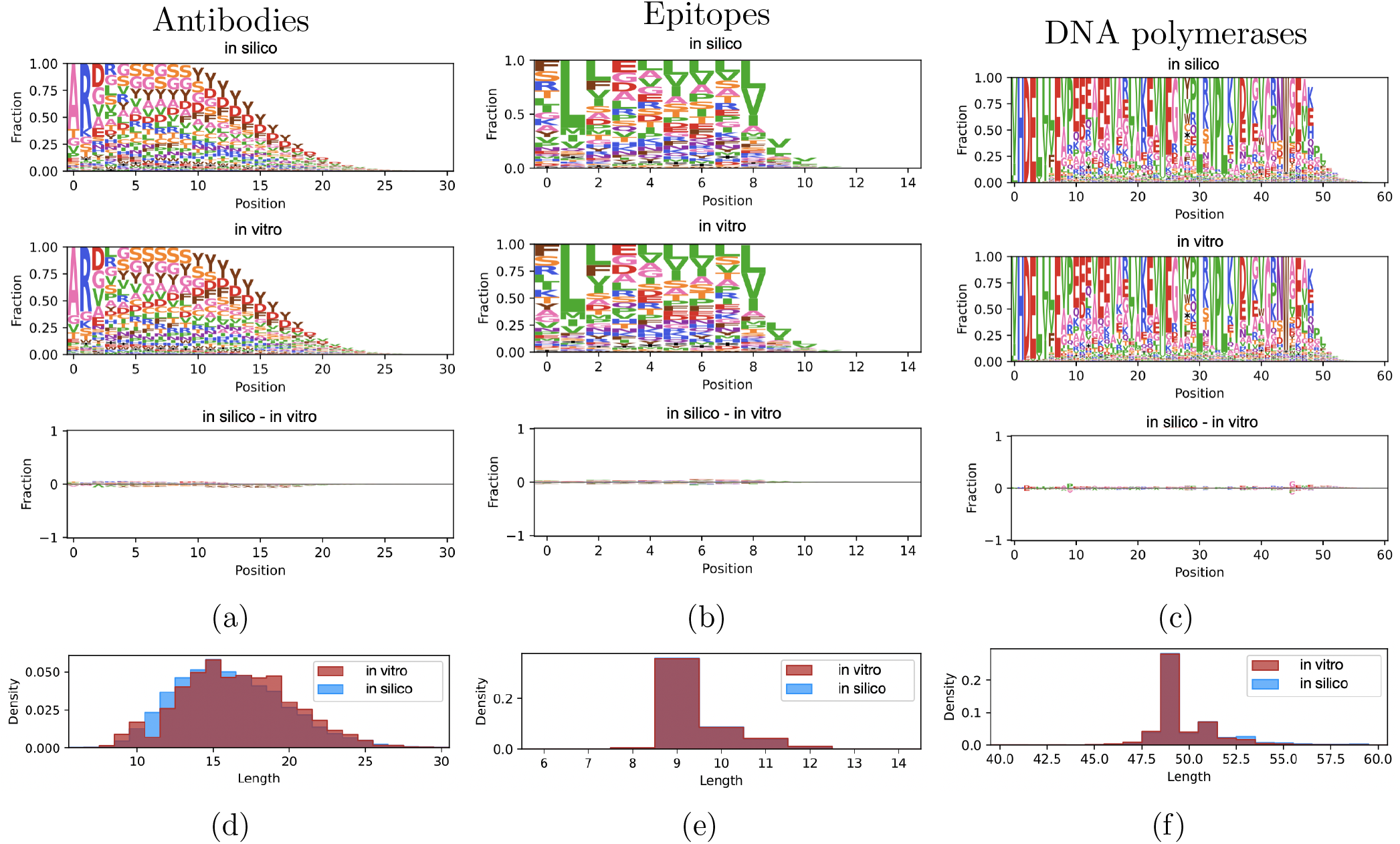
Comparing amino-acid usage and length distributions in the *in silico* and *in vitro* variational synthesis libraries. **a-c**, Each logo in the first two rows shows the marginal distribution over amino acids. Formally, in the notation of Appendix C.1, each position plots 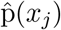, the estimated marginal distribution over amino acids (with the padding character $ from shorter sequences represented as white space). The third row plots, as a logo, the difference between the marginal distributions, 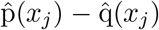, where 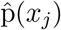 comes from *in silico* samples and 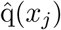 comes from *in vitro* manufactured samples. **d-f** Distribution of sequence lengths. **a**,**d** Antibody dataset. **b**,**e** Epitope dataset. **c**,**f** DNA polymerase dataset.

**Figure S2:**
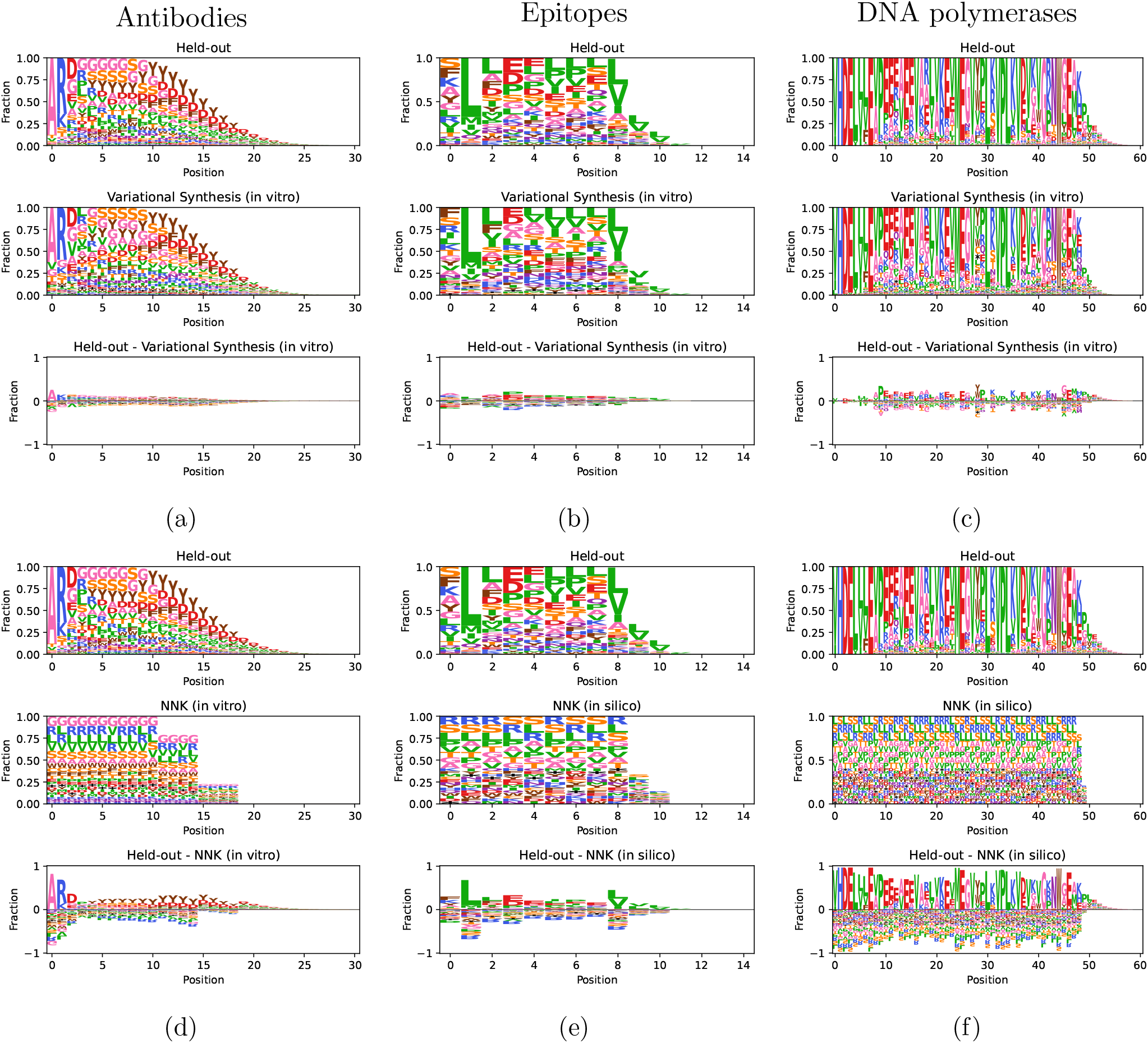
Comparing amino-acid usage in the held-out data and the *in vitro* manufactured variational synthesis library. **a-c**, Same as Figure S1a-c, except comparing held-out data to the manufactured variational synthesis library. **d-f**, Same as a-c, except comparing held-out data to *in silico* samples from the manufactured NNK library (d) and *in silico* NNK library (e, f).

**Figure S3:**
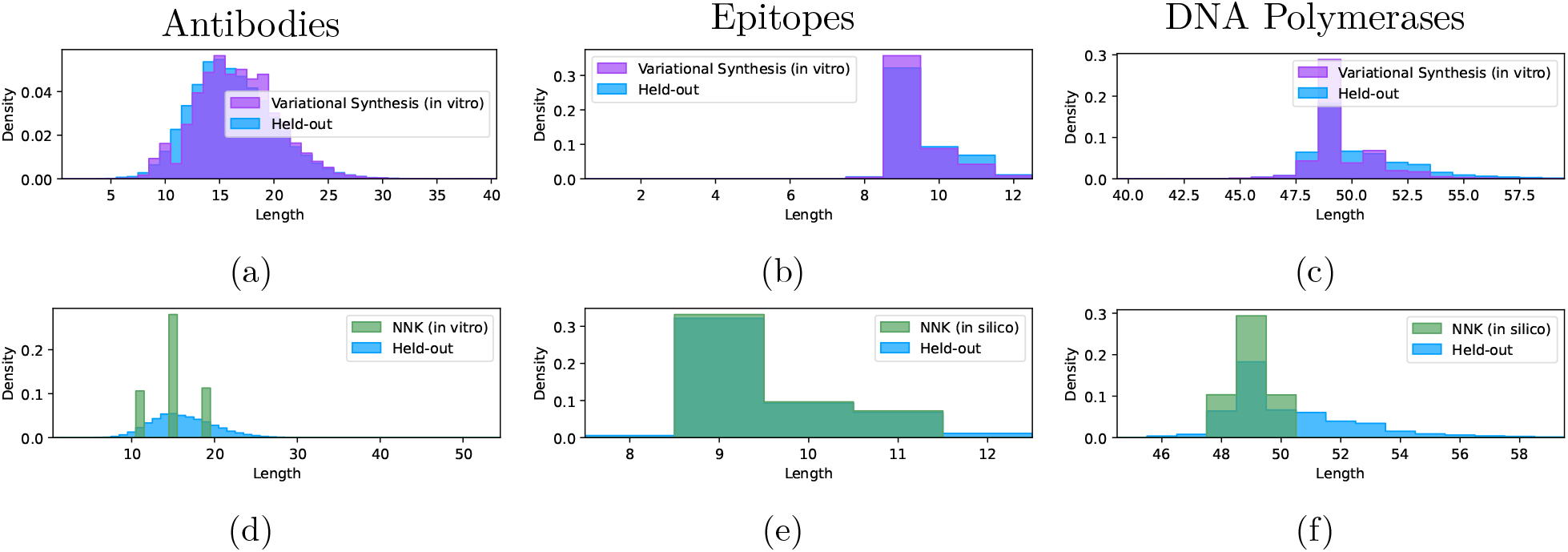
Comparing length distributions in the held-out data versus the *in vitro* variational synthesis libraries and versus the NNK libraries. **a-c**, Same as Figure S1d-e, except comparing held-out data to the manufactured variational synthesis library. **d-f**, Same as a-c, except comparing held-out data to samples from the *in vitro* manufactured NNK library (d) and *in silico* NNK library (e, f).

### C.3 Nonparametric tests (BEAR and MMD)

#### Setup

The basic goal of generative modeling is to develop an estimate p_*θ*_^*⋆*^ (*x*) of the distribution p_0_(*x*) underlying the data, from which we can draw new samples. In particular, the aim is for samples *X ∼* p_*θ*_^*⋆*^ (*x*) from the model to be indistinguishable from samples *X ∼* p_0_(*x*) from the data distribution.

A variety of metrics have been developed to evaluate generative biological sequence models, with different authors focusing on different desiderata. Though the details vary, many of these evaluation procedures can be described as special cases of a general procedure. They start with a function of sequences *F*(*·*) : *χ → ℝ*^*d*^, which outputs a scalar or vector summary of the sequence’s features. A simple example is *F*(*x*) = *x*_*i*_, which outputs a one-hot encoding of the amino acid at position *i*. ^23^ A more complex example is *F*(*x*) = pLLDT(*x*), the prediction confidence of a protein folding algorithm (ESMFold) applied to *x*. ^17^ More broadly, *F*(*x*) can be any hand-crafted feature of *x* or the output of a machine learning model such as a supervised predictor or unsupervised embedding. Once *F*(*x*) is chosen, it can be used to compare samples from the generative model to (held-out) samples from the data, e.g. by examining the error

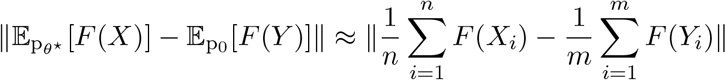

where *X*_1:*n*_ *∼* p_*θ*_^*⋆*^ (*x*) are samples from the model and *Y*_1:*m*_ *∼* p_0_(*x*) are data. In sum, typical generative model evaluations check that samples from the model look like real data according to a chosen metric *F*(*x*). Indeed, our analysis of average amino acid usage, length distribution, and low-dimensional representations can each be seen as instantiations of this general idea. (Note also that popular metrics for generative image models, such as the Fr’echet Inception Distance and the Inception Score, can also be framed in a similar way. ^24;5^)

These approaches are limited, however, because *F*(*x*) can only capture certain aspects of the data. Even if the model samples look similar to the data according to chosen metrics, they may differ dramatically along other dimensions. Adding additional metrics, i.e. alternative values of *F*(*x*), can reduce this problem but does not eliminate it. Moreover, since some generative models are optimized for certain metrics and not others, performance can very widely among different models, and results are incomparable.

To overcome these limitations and pursue a more unbiased evaluation, we used nonparametric two-sample tests. Rather than focus on specific features of sequences, these methods evaluate how well the entire model distribution p_*θ*_^*⋆*^ (*x*) matches the data distribution p_0_(*x*). To accomplish this, they implicitly consider *all* possible summary functions *F*(*x*), making use of an infinite number of features. Hence, they resolve the problem of relying on a finite set of hand-picked metrics *F*(*x*).

Nonparametric tests provide more global judgments of generative model quality than handpicked features, but different nonparametric tests can still place different emphasis on different features of the data. This can result in different conclusions in practice, when the tests are applied to finite datasets. To mitigate this issue, we make use of two different nonparametric two-sample tests, both designed for biological sequence data. The BEAR test is based on an autoregressive model based on variable-length kmers, while the MMD uses a kernel involving the Hamming distance between sequences. As a result, the BEAR test places somewhat greater focus on local correlations between nearby amino acids, while the MMD considers more global structure.

To make the computation more feasible, we ran each test on a subsample of sequences drawn uniformly at random from the model-generated sequences and the data-generated sequences. We used 200,000 sequences form each in our subsample.

#### MMD

First, we used maximum mean discrepancy to evaluate the mismatch between model and data. ^4^ It is defined as

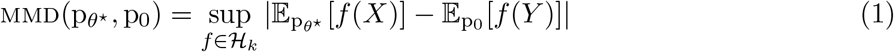

where *ℋ*_*k*_ is a reproducing kernel Hilbert space with kernel *k*(*x, y*). The space *ℋ*_*k*_ can contain an infinite number of functions, allowing the MMD to check all possible directions in which the model and data distributions may differ. We can think of each *f ∈ ℋ*_*k*_ as a possible sequence-to-phenotype mapping, so the MMD can be interpreted as the worst-case difference in average phenotype between the two distributions.

The MMD can be estimated using samples from the model samples *X*_1:*n*_ *∼* p_*θ*_^*⋆*^ (*x*) and held-out data *Y*_1:*m*_ *∼* p_0_(*x*) as

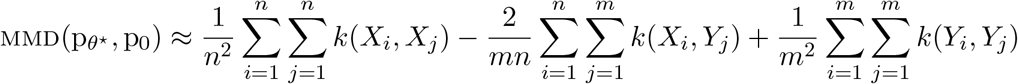

(We use the V-statistic rather than the U-statistic, which ensures the estimated MMD is greater than zero.) We report this value as a measure of model-data mismatch in Figure 2g and elsewhere in the main text.

In practice, we use the heavy-tailed Hamming kernel,

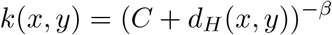

where *β* = 0.5, *C* = 1, and *d*_*H*_ is the Hamming distance, i.e. the number of mismatches between *x* and *y*. We use the Hamming distance because the number of mutations between two sequences is a common and often reasonably accurate way of evaluating how functionally similar two sequences are likely to be. We chose *β <* 1 to allow for a lower penalty between distant sequences, avoiding “diagonal dominance”. ^3^ We padded the sequences to the same length on the right with stop symbols ($), so that when *x* and *y* have different length, the difference in lengths is added to the Hamming distance. Using this kernel, the MMD is guaranteed to be greater than zero if and only if p_*θ*_^*⋆*^ ≠ p_0_. Formally, this is guaranteed by the fact that the kernel has discrete masses, and hence is characteristic (Example 6 in Amin et al. (2023)^3^).

#### BEAR

We applied the BEAR two-sample test proposed in Amin et al. (2021). ^1^ It is a nonparametric Bayesian test, which computes a Bayes factor comparing the hypothesis that the model does not match the data to the hypothesis that it does. That is, it compares the hypothesis p_*θ*_^*⋆*^≠ p_0_ to p_*θ*_^*⋆*^ = p_0_. Given model samples *X*_1:*n*_ *∼* p_*θ*_^*⋆*^ (*x*) and held-out data *Y*_1:*m*_ *∼* p_0_(*x*), the Bayes factor is

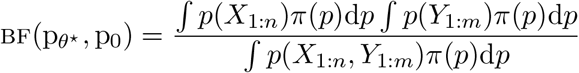

Here, *π*(*p*) is a prior distribution that covers all possible sequence distributions, namely the BEAR model distribution. The denominator computes the likelihood of the data assuming *X*_1:*n*_ and *Y*_1:*n*_ come from the same distribution, integrating over all possible distributions. The numerator computes the likelihood of the data assuming *X*_1:*n*_ and *Y*_1:*n*_ come from different distributions, again integrating over all possible distributions. The ratio is the odds in favor of the hypothesis that p_*θ*_^*⋆*^≠ p_0_.

In detail, we set the hyperparameters of the BEAR test as follows. First, we set the maximum considered kmer length to the maximum sequence length in the dataset (which has very small posterior probability in practice compared to shorter sequences). Second, for convenience, we used a uniform autoregressive base model and a Jeffreys prior on the transition probability, corresponding to *f*(*θ*) = 1 and *h* = 2 in the notation of Amin et al. (2021). ^1^

To make the Bayes factor more interpretable and comparable across models and datasets, we report the geometric mean Bayes factor per sequence 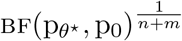 in Figure 2g, Figure 3g and Figure 4g. We use *n* = *m* in practice, so model and data contribute equally. Intuitively, this average Bayes factor can be interpreted as the evidence provided by each sequence that the model and data distributions are distinct.

### C.4 Witness function

To estimate the fraction of *in silico* or *in vitro* designs that are realistic, we used the witness function of the MMD. ^13^ The witness function is the function with the maximum difference in expected value under the model and data distributions, i.e. the function that attains the supremum in Equation (1). It is given by

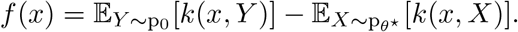

It can be estimated empirically from model samples *X*_1:*n*_*∼* p_*θ*_^*⋆*^ (*x*) and held-out data *Y*_1:*m*_*∼* p_0_(*x*) as,

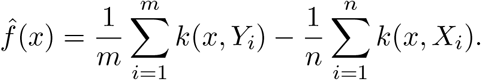

The witness function measures how much a sequence *x* looks like it came from the data distribution p_0_(*x*) as compared to the model p_*θ*_^*⋆*^ (*x*). Sequences that look like outliers with respect to the data will receive low scores according to the witness function. Sequences that are heavily over-represented in the model samples, as compared to the data, will also receive low witness function scores. Note that as a result, the witness function takes into account some notion of diversity as well as realism, and will not overly favor models that just produce a limited set of points that are inliers for the data distribution. This distinguishes the witness function from other methods for outlier detection, and makes it particularly appropriate for evaluating generative models.

We use the witness function to evaluate the realism of our generated samples. Specifically, let 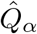 denote the *α*-quantile of the empirical distribution of witness function scores,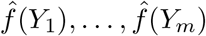.Then, we compute the fraction of “clearly realistic” sequences as the fraction of generated sequences above this threshold,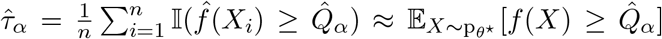. If the model closely matched the data distribution, we would expect that the fraction of sequences above 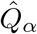 would be 1 *− α* on expectation. In practice it is lower, due to model-data mismatch. We report a threshold of *α* = 0.33 in the main text, but we also show 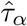 for all values of *α* in the supplementary figures (Figure S5b, Figure S6).

### C.5 NNK Libraries

To benchmark variational synthesis against conventional high-diversity library synthesis techniques, we compare to NNK libraries. In an NNK library each codon is generated randomly, such that the first two positions (N) can be any base while the final position (K) is a G or T. To help evaluate NNK libraries, we specify the distribution of sequences it produces formally, using a synthesis model. We work in the setup and notation of Weinstein et al. (2022). ^32^

Typically NNK libraries are used with one or a few different lengths. Let ℒ ⊆ {1, 2, …} be the set of lengths, and *v*^*L*^ a vector describing their relative concentrations. Then we have the generative model,

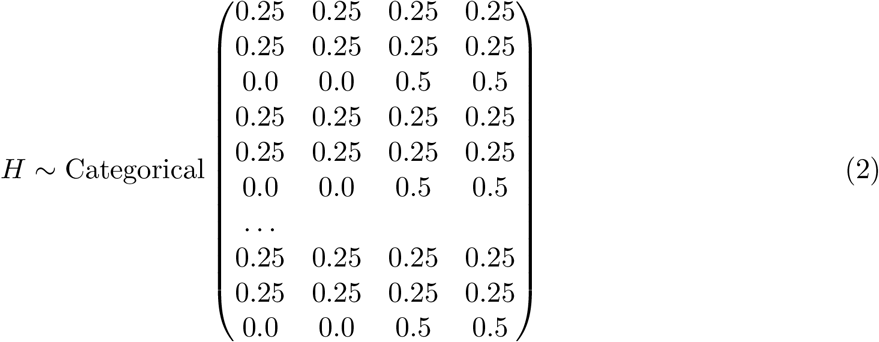

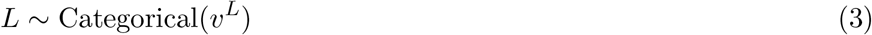

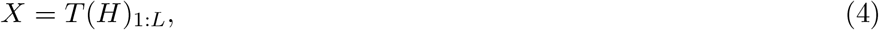

which can be interpreted as follows. Equation (2) describes a distribution over DNA, with each column of the matrix giving the probability of A, C, G and T respectively. Each nucleotide is sampled independently according to these probabilities. Equation (3) samples the length of the sequences, choosing from among the lengths in ℒ according to their relative concentrations *v*^*L*^. Equation (4) translates the DNA sequences into protein: the function *T*(*·*) maps DNA sequences to amino acid sequences according to the standard genetic code. The final result *X* is a sample amino acid sequence. Note we can also calculate the likelihood of data under this model, e.g. using the tools in Weinstein et al. (2022). ^32^

Specifically, we compared our variational synthesis libraries to the following NNK libraries, with lengths chosen based on typical values in each training dataset (there is no established procedure for choosing lengths in degenerate codon libraries except to match the typical lengths of real sequences). We match the relative concentration *v*^*L*^ to the relative frequency of each of the lengths in ℒ in the training data.

1. Antibodies: We created an *in vitro* library with lengths ℒ = {11, 15, 19}. For evaluations, we sequenced a random subset of the library: 4,494,385 sequences after assembly and filtering (Appendix A).
2. Epitopes: We drew samples *in silico* from the generative NNK library model above, with ℒ = {9, 10, 11}.
3. DNA Polymerases: We drew samples *in silico* from the generative NNK library model above, with ℒ = {48, 49, 50}.

**Figure S4:**
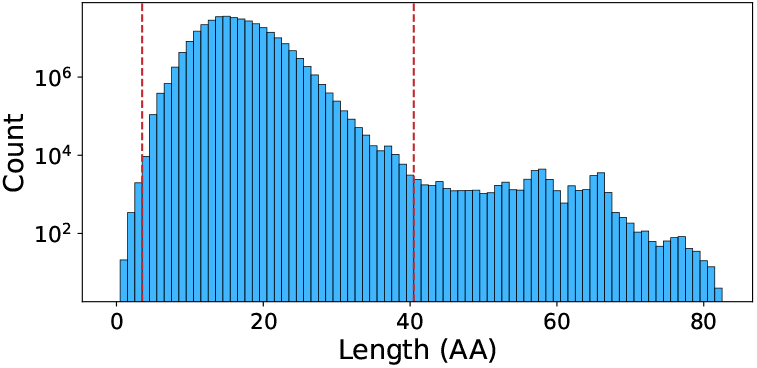
Length distribution in the antibody OAS dataset. Sequences with lengths between the vertical lines were included in the training and held-out data for variational synthesis. Note the y-axis is on a log scale.

To compute finite perplexities describing how well these models match real sequence data, we use only data sequences with lengths in *L*. Hence the perplexities we report should be taken as a conservative estimate of how well the NNK libraries match the real data; in fact they perform worse than the perplexity would suggest, since they do not match the length distribution.

## D Details on antibodies

In this section we provide details on our procedure for synthesizing CDRH3 sequences.

### D.1 Dataset

We train our synthesis model on the human CDRH3 sequences from the Observed Antibody Space^19^ database. We collected all human repertoires available in the database on 2024-06-07: 13,265 repertoires total, from 756 donors. Then, we excluded all repertoires labeled with diseases that may lead to production of potentially harmful antibodies: the full list in the OAS nomenclature is ‘Healthy/celiac-disease’, ‘Allergy/NoSIT’, ‘Allergy/SIT’, ‘MS’, ‘MuSK-MG’, ‘AChR-MG’, ‘SLE’, ‘Light-Chain-Amyloidosis’, ‘Asthma’, ‘Allergic-Rhinitis-Out-Of-Season’, ‘Allergic-Rhinitis-In-Season’. This left us with 11,271 repertoires from individuals aged 3 to 92 (42 *±*19) that contained 327,341,295 unique productive CDR3 amino acid sequences (CDR3 was determined as in OAS, i.e., positions 105 to 117 in the IMGT ^12^ numbering scheme). We evaluated the distribution of lengths of these sequences (Figure S4) and excluded extreme cases – sequences with length below 4 or above 40, which resulted in 325,596,608 unique sequences. We then uniformly split this set of sequences into 90% training and 10% held-out data.

### D.2 Antibody language model

We used a state-of-the-art generative antibody language model IgLM ^27^ to generate sequences *in silico*. IgLM was trained on the OAS database as well, although in this case, the authors used full-length sequences from all species and chains, and added chain and species tokens to the model input. We generated 200 000 full-length heavy chain human antibody sequences using tokens [HEAVY] and [HUMAN], and then extracted CDR3s from the resulting sequences. Note that it is not possible to evaluate likelihoods of arbitrary CDR3 sequences using IgLM since it was trained to generate or infill full-length sequences.

**Figure S5:**
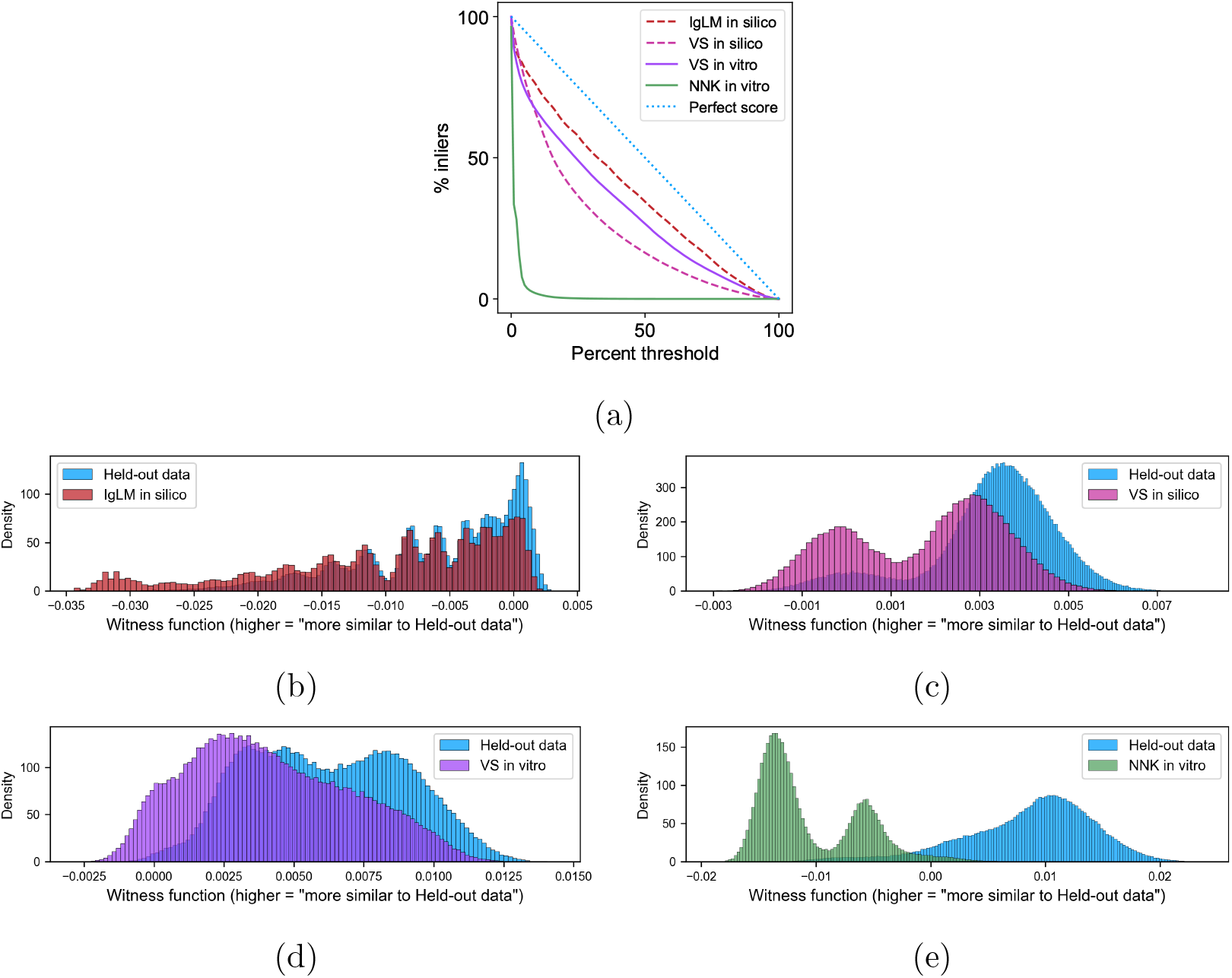
Witness function analysis of antibodies. **a**, % of generated sequences (*in silico* or *in vitro*) with witness function values above the real-data percentile threshold given on the x-axis (VS: variational synthesis). Formally, this is a plot of 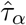 as a function of *α*(Appendix C.4). **b**, Distribution of witness function scores 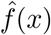 for *in silico* samples from IgLM versus held-out data. **c**, Distribution of witness function scores for variational synthesis *in silico* samples versus held-out data. **d**, Distribution of witness function scores for variational synthesis *in vitro* manufactured samples versus held-out data. **e**, Distribution of witness function scores for NNK *in vitro* manufactured sequences versus held-out data.

**Figure S6:**
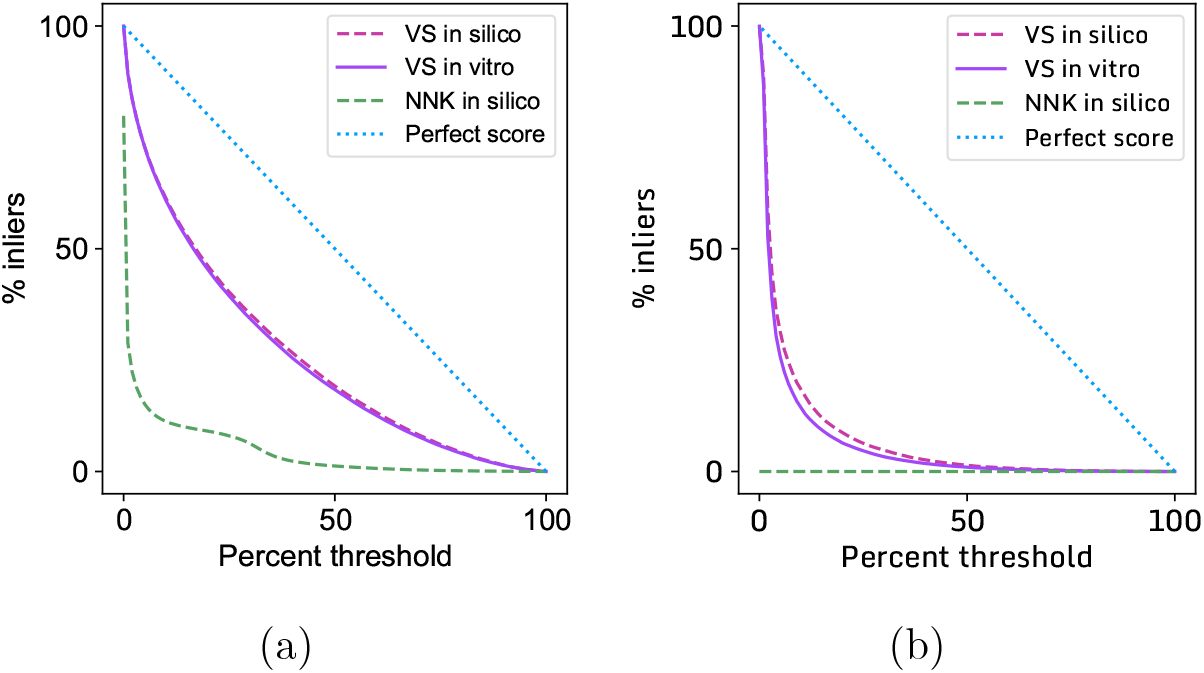
Witness function analysis of epitopes and DNA polymerases. Each plot shows the % of generated sequences (*in silico* or *in vitro*) with witness function values above the realdata percentile threshold given on the x-axis. Formally, these are plots of 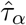 as a function of *α*(Appendix C.4). **a**, Results for epitope models and libraries. **b**, Results for DNA polymerase models and libraries.

**Table S2:** Percentage of sequences in each library that are predicted to bind HLA-A*02:01 by NetMHCpan.

|  | Non-binders | Weak binders | Strong binders |
| --- | --- | --- | --- |
| Training set | 0.0% | 0.0% | 100.0% |
| NNK (*in silico*) | 97.3% | 2.0% | 0.7% |
| VS (*in silico*) | 39.2% | 25.5% | 35.4% |
| VS (manufactured) | 38.2% | 25.5% | 36.3% |

## E Details on HLA-A*02:01 epitopes

We train our synthesis model on the results of a virtual screen for HLA-A*02:01 binders. The screen selects sequences that are predicted to be strong binders by NetMHCpan, version 4.1. ^22^ We can formally describe this data as samples from a conditional generative model of peptides.

Let the function *b*(*·*) : *𝒳 →* {0, 1} denote the NetMHCpan model for HLA-A*02:01, which takes in an amino acid sequence *x* and outputs a prediction for whether it is a strong binder for HLA-A*02:01 (*b*(*x*) = 1) or not (*b*(*x*) = 0).

We consider a global natural distribution over possible antigens *q*(*x*). In particular, let *v* be length 20 vector (on the simplex) denoting the frequency of each amino acid across proteins. We use the amino acid frequencies for the human proteome ^25^, but similar frequencies hold across evolution (e.g. see Fig. S3 of^21^). Then, 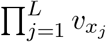 is the distribution over sequences where *L* amino acids are each sampled independently from *v*. Now we define the global distribution over possible antigens as a uniform mixture across the most frequent lengths for class I HLAs (8-12 amino acids) ^29^:

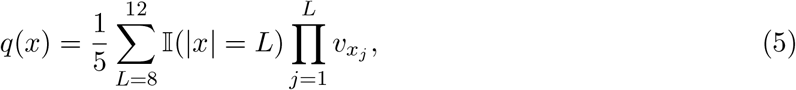

where 𝕀(|*x*| = *L*) takes value one when *x* has length *L*. Note although *q*(*x*) is a simple generative model, which ignores correlation structure among nearby amino acids, we have observed in practice that it provides a reasonable approximation of plausible antigen distributions: there are a vast array of possible antigens (including human, bacterial, and viral proteins), and at short length scales there is little common structure among their sequences.

Our goal is to synthesize antigens presented on HLA-A*02:01. We therefore aim to synthesize samples from the conditional generative model describing the distribution of antigens predicted to bind HLA-A*02:01,

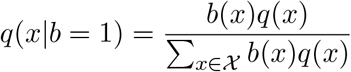

We can draw samples from this distribution via rejection sampling: draw a sample *X ∼ q*(*x*) and accept it if *b*(*X*) = 1, otherwise reject and repeat. This rejection sampling procedure is, precisely, a virtual screen of samples from *q*(*x*) using NetMHCpan. Note we can also estimate the normalizing constant Σ_*x∈𝒳*_ *b*(*x*)*q*(*x*) = 𝔼_*q*_[*b*(*X*)] as the fraction of samples that are accepted. This allows us to estimate the likelihood *q*(*x*|*b* = 1) of the true distribution, and hence compute its perplexity.

Finally, we train our synthesis model on samples from *q*(*x b* = 1) and run the corresponding synthesis procedure.^1^

## F Details on DNA polymerases

We train our synthesis model on samples from a generative evolutionary model, conditioned on an initial segment of the Taq polymerase. In particular, let *q*(*x*) denote the distribution of Progen2 (`progen2-xlarge`), a protein language model trained on a large and diverse dataset (including UniRef and the BFD metagenomic dataset). We prompt the model with the first *𝓁* = 782 amino acids of Taq (Uniprot ID `DPO1_THEAQ`), removing the last 50 amino acids. The conditional distribution of the model can then be written *q*(*x* | *x*_1:782_ = taq_1:782_). This is a prediction of proteins likely to be generated by evolution, given that their initial segment matches *Taq*. We can draw samples from this distribution by generating from the model with the temperature parameter and nucleus sampling probability parameter both set to one.

In practice we found that many generated samples were quite long, in some cases with no stop codon at all. To ensure oligosynthesis was tractable without assembly, we set a length cutoff of *T* = 60 amino acids, rejecting samples from the model that go beyond this length before they reach a stop codon. This gives us a sample from *q*(*x* | *x*_1:782_ = taq_1:782_, |*x*| *−* 782 *≤* 60). We take this as our target distribution, and train our synthesis model on samples from this distribution. Note our analysis of these samples focuses just on the completion, ignoring the constant region.

To compute the likelihood and perplexity of sequences under this distribution, we use Bayes’ rule,

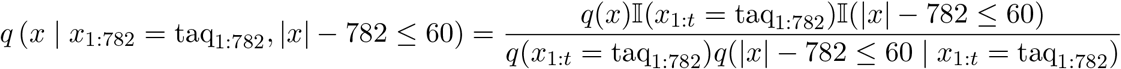

where 𝕀(*·*) is the indicator function which takes value 1 if the statement is true and 0 otherwise. The likelihood *q*(*x*) can be evaluated for ProGen2 since it is an autoregressive model.^2^ The term *q*(*x*_1:*t*_ = taq_1:782_) can be evaluated similarly. Finally, *q*(|*x*| *−* 782 *≤* 60 | *x*_1:*t*_ = taq_1:782_) can be estimated as the fraction of samples that were accepted during rejection sampling.

## Footnotes

1 In practice our training procedure removes duplicated samples, but since only 0.06% of the samples drawn from *q*(*x*|*b* = 1) appear more than once, the difference between the de-duplicated distribution and *q*(*x*|*b* = 1) is minimal.

2 Note computing *q*(*x*) requires a small modification to the likelihood computation code provided with the ProGen2 model: we just care about the likelihood in the forward (i.e. N terminal to C terminal) direction, not the reverse direction, and the stop symbol must be included since it is part of the generative process

